# AFFIPred: AlphaFold2 Structure-based Functional Impact Prediction of Missense Variations

**DOI:** 10.1101/2024.05.13.593840

**Authors:** Mustafa Samet Pir, Emel Timucin

**Affiliations:** Acibadem University, Institute of Health Sciences, Department of Biostatistics and Bioinformatics, Atasehir Istanbul 34752 Turkey; Acibadem University, School of Medicine, Department of Biostatistics and Medical Informatics, Atasehir Istanbul 34752 Turkey

## Abstract

Structural information holds immense potential for pathogenicity prediction of missense variations, albeit structure-based pathogenicity classifiers are limited compared to their sequence-based counterparts due to the well-known gap between sequence and structure data. Leveraging the highly accurate protein structure prediction method, AlphaFold2 (AF2), we introduce AFFIPred, an ensemble machine learning classifier that combines established sequence and AF2-based structural characteristics to predict disease-causing missense variant pathogenicity. Based on the assessments on unseen datasets, AFFIPred reached a comparable level of performance with the state-of-the-art predictors such as AlphaMissense and Rhapsody. We also showed that the recruitment of AF2 structures that are full-length and represent the unbound states ensures more precise SASA calculations compared to the recruitment of experimental structures. Second, in line with the the completeness of the AF2 structures, their use provide a more comprehensive view of the structural characteristics of the missense variation datasets by capturing all variants. AFFIPred maintains high-level accuracy without the well-known limitations of structure-based pathogenicity classifiers, paving the way for the development of more sophisticated structure-based methods without PDB dependence. AFFIPred has predicted over 210 million variations of the human proteome, which are accessible at https://affipred.timucinlab.com/.

## Introduction

Significant advances have been made in computational methodologies for the classification of the functional impact of missense variants. Pioneer classifiers mainly relied on sequence-based attributes, such as conservation scores that could be obtained from homolog protein sequences and their multiple sequence alignments [1, 2]. Notwithstanding the undeniable power of sequence-based attributes, structural information has also been shown to contribute to pathogenicity classification of missense variations [3–7].

Structure-based classifiers mostly recruit static protein structures to account for certain structural characteristics of the variant in question, while a few methods additionally leverage protein dynamics that could be accessed by all-atom or coarse-grained molecular dynamics (MD) simulations to address the functional impact of variants [8, 9]. Solvent accessible surface area (SASA) and mean square fluctuation (MSF) are among the notable structural attributes that could be calculated using static structures [10] and classical MD approaches [8, 11, 12], respectively. Despite the apparent contribution of structural attributes to the classification task, only a limited number of structure-based pathogenicity predictors are available.

The limited usefulness of protein structures in pathogenicity classification has historically stemmed from the gap between available sequence data and known protein structures [8, 13, 14]. However, this gap has narrowed significantly due to highly accurate protein structure prediction methods. Essentially, the artificial intelligence system, AlphaFold2 (AF2), being the pioneer method [15] produced strikingly accurate predictions in the CASP14, surpassing all other methods [16]. Currently, more than 200 million AF2-computed protein structures are deposited in the AlphaFold Protein Structure Database (https://alphafold.ebi.ac.uk/) [17, 18], including the human proteome [19]. Furthermore, the RCSB Protein Data Bank (RCSB PDB) currently accommodates more than 1 million AF2 computed structural models. Unequivocally, these transformative developments in the field of protein structure prediction have reshaped and greatly expanded protein structure databases [20, 21], presenting a valuable opportunity for structure-based pathogenicity classifiers to expand their applicability.

While PDB offers valuable structural information for structure-based classifiers, PDB entries often lack complete structures, especially for large proteins and/or intrinsically disordered regions (IDRs) [22, 23]. Excluding these variations from unstructured regions from the training data of structure-based classifiers can further introduce bias, limiting the usefulness of protein structure for pathogenicity prediction. AF2-computed structures that provides a complete structural characterization of a given protein along with a built-in confidence score [16] may offer an advantage at this point. Accordingly, the place of AF2 structures in pathogenicity classification has recently been explored. Schmidt et al. (2023) demonstrated the use of AF2 structures to extract detailed structural information for classifying pathogenicity [7]. Similarly, the AlphaMissense method, built on the original AlphaFold system and trained on a large variant database, also uses the AF2 structures to predict the pathogenicity of missense variants [24]. Both studies highlight the promise of AF2-derived structures for understanding pathogenicity, implying the capacity of AF2-derived structures to replace experimental structures in pathogenicity prediction tasks. Despite these successful examples, there is a lack of AF2-based structure classifiers that harmonize both sequence and structural information to address variant pathogenicity. Furthermore, a critical question still remains: can AF2 structures effectively substitute existing PDB structures? This is particularly critical for proteins containing IDRs. As such, predictions of IDRs based on AF2 may be unreliable, and the absence of corresponding structures in the PDB may further prevent their validation.

Here, we built ensemble machine learning models that combine sequence- and AF2-based structural features to classify disease-causing missense variants. Our models, named AFFIPred, were trained on a large dataset of missense variations collected from ClinVar. Our results showed that the combination of AF2-based structural features with well-recognized sequence-based features improved the predictions of many sequence or structure-based predictors without reliance on experimental structures. Overall, our findings reflect the complementarity, sometimes superiority, of AF2 structures to their experimental counterparts in pathogenicity classification of missense variations.

## Materials and methods

### Dataset Collection

We collected all human missense variations submitted to the ClinVar database [25] as of April 2023. Variations with at least one review star were selected and matched to entries in the UniProt Knowledge Base [26]. Duplicate variations sharing the same genomic coordinates were eliminated. Subsequently, various sequence and structural features were computed for each variation, discarding those for which any of the feature computation failed.

Among sequence-based features, we initially collected the number of Gene Onthology (GO) identifiers for each protein. Position-specific independent count (PSIC) scores were obtained from the pre-computed profiles of PolyPhen-2 [3, 27, 28]. Hydrophobicity and volume of the native and mutant positions were calculated using the Kyte-Doolittle scale [29] and amino acid volume measurements from the ref. [30], respectively. BLOSUM62 matrix was used to extract the substitution scores [31]. Structural features were computed using the AF2 computed structures (v4), which were obtained as a part of the human proteome (UP000005640) of AlphaFold-EBI structure DB (https://alphafold.ebi.ac.uk) [15, 17].

Selection of AF2 structure is carried out based on the variant position. If the variant position is equal or higher than 2701, the AF2 structure was selected from overlapping AF2 models to cover at least 200 aa upstream and 200 aa downstream of the position. Otherwise, i.e. the variant position was less than 2701, the F1 model of AF2 structure was utilized for computation of the structural attributes. As an example, for the variant positions of the KMT2D protein (Uniprot: O14686) falling between 3201 and 4200, the F16 model encompassing positions 3000 to 4400 was used.

The selected AF2 structure was used to determine the residue confidence score (pLDDT), and relative accessibility surface area (rASA) for the variations using the dssp module of BioPhyton.PDB [32–35]. We also computed the central tendency and spread measures of the pLDDT score distribution of the selected AF2 models. The resulting dataset is available at the https://github.com/timucinlab/AFFIPred.

### Model Construction

An extreme gradient boosting (XGBoost) algorithm was employed, which is an ensemble learning technique based on the gradient-boosted decision-tree method and known for its capability in handling large datasets [36]. Hyperparameter tuning was performed using the Hyperopt framework [37], a Bayesian optimization library known for its efficiency in searching the hyperparameter space. Two hundred evaluations were conducted to find the optimal combination of hyperparameters, maximizing the evaluation metric of AUROC (AUC). The optimized parameters and tested ranges for each parameter were given in S1. Three distinct models were trained employing a nested cross-validation strategy with an outer loop of 10 folds and inner loop of 5 folds for the hyperparameter tuning, model selection and error estimation.

### Performance Assessment

Three distinct variant sets that were unseen to each AFFIPred model was used for evaluation. The proteins that constitute the entire Clinvar dataset was split into three equal parts. The variations of each protein subset was separated as an unseen dataset While the rest of data was used for the training. Performance of AFFIPred was evaluated against 36 methods from ANNOVAR [38], 3 additional methods, namely Rhapsody, EVmutation, and Polyphen-2, AlpScore [7] and AlphaMissense [24] were compared against AFFIPred. The predictions for EVmutation and Polyphen-2 were obtained through Rhapsody [6, 8]. We evaluated the classification performance using the metrics of AUC, F1 score, MCC, accuracy, and Spearman’s rank correlation coefficient.

### Prediction of Missense Variations in the Human Proteome

The AF2 prediction (v4) of the human proteome (UP000005640), composed of 20,597 proteins, was retrieved from the AlphaFold-EBI structure DB (https://alphafold.ebi.ac.uk) [15, 17]. All possible variations of the proteins in the human proteome was generated and their sequence and structure-based features were computed using the AF2 structures of each protein. Selection of AF2 structures was done similarly described for the ClinVar set. All variations were predicted by each AFFIPred classifier. Pathogenicity labels were assigned based on the mean probability scores of three predictions. As such, the variant is classified as neutral if its mean probability score is less than 0.5, while it is denoted as pathogenic if the score is equal or higher than 0.5. We have developed an R Shiny based web-interface, wherein all the possible variants in the human proteome were pre-calculated. Any single amino acid variation can be searched using amino acid position, genomic position and dbSNP id. These predictions can be accessed through https://affipred.timucinlab.com/. We have also developed a command line tool to facilitate handling of high number of queries and easy integration with pipelines.

## Results and Discussion

### ClinVar Collection of Missense Variations and Their PDB Status

We compiled all of the missense variations that had at least one reviewer star rating in the ClinVar as of April 2023. While the initial collection included over 90,000 variants, only 75,330 could be processed during feature calculation, resulting in a refined, non-redundant dataset of missense variants from ClinVar. This variation dataset has been composed of 7,813 distinct proteins. 54% of the proteins that were associated with an at least one PDB entry were labeled as the PDB group, while the remaining 46% without any experimental structures were labeled as the NA group (Fig. 1a). The experimental methods used for the structure determination of the PDB group were dominated by X-ray crystallization among other techniques such as nuclear magnetic resonance and electron microscopy (Fig. 1a), matching with the overall PDB statistics.

**Fig 1.**
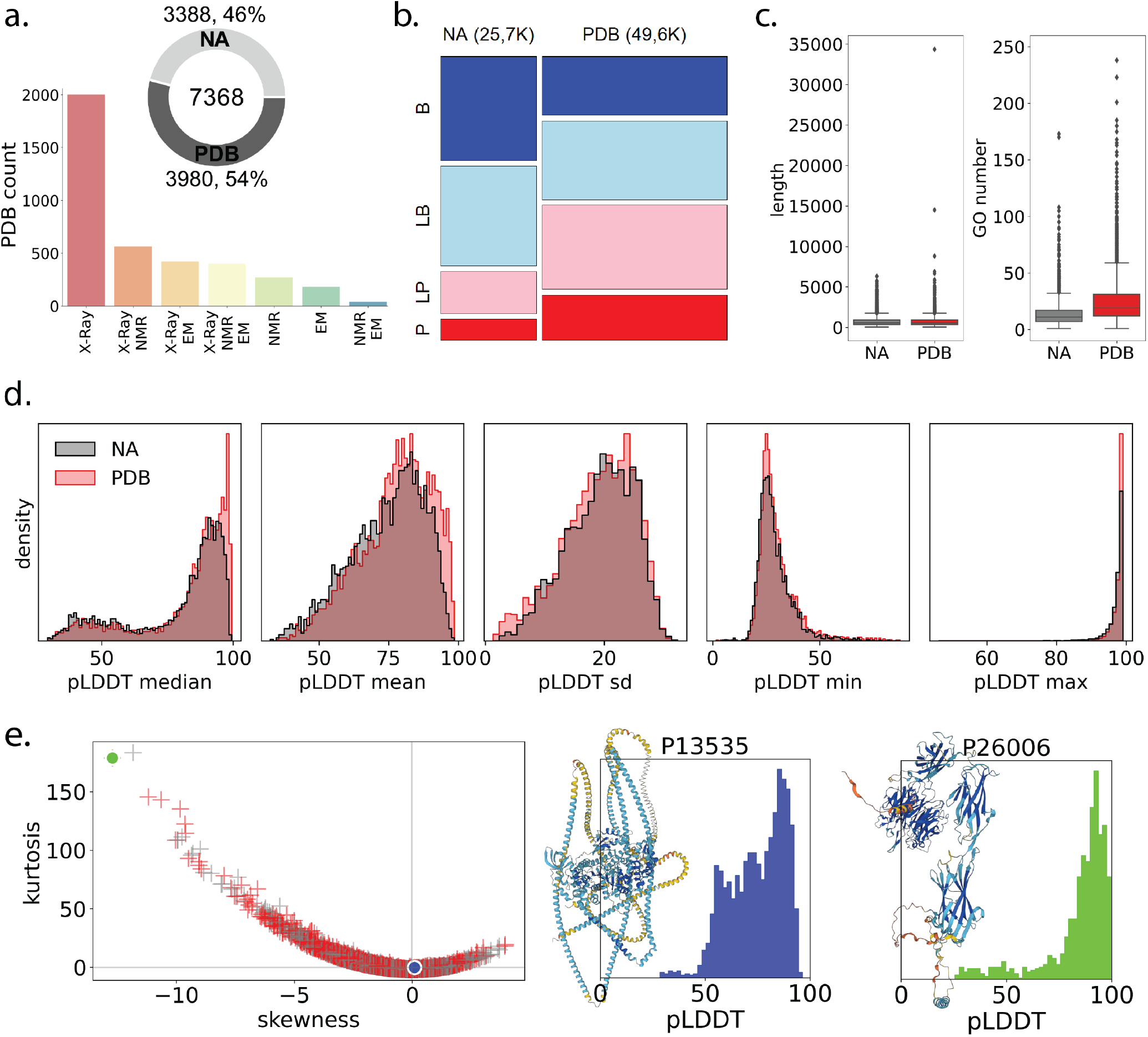
ClinVar Collection of Missense Variations and Associated Structural Data. (a) Pie chart shows the number of unique proteins that are linked to at least one PDB entry (PDB) and that are not linked (NA). Bar chart shows how often different experimental methods were used to determine the 3D structure of proteins in the PDB group. (b) Mosaic plot visualizes how the distribution of pathogenicity differs between PDB and NA protein groups (B: benign, LB: likely benign, LP: likely pathogenic, P: pathogenic). (c) Box plot compares the number of Gene Ontology (GO) terms associated with proteins in the PDB and NA groups. Plots in (d) show the statistics of pLDDT distributions of the AF2 structures of 7368 proteins. Left plot in (e) Skewness-kurtosis plot displays the pLDDT score distributions. Proteins from the PDB group are marked with red crosses, while those from the NA group are marked with grey crosses. Two representative distributions for the Uniprot IDs of P13535 and P26006 were plotted. The former protein has a pLDDT distribution that approximates a normal curve with skewness and excess kurtosis values close to zero while the latter distribution is leptokurtic and negatively skewed. AF2 structures of both structures were visualized next to each plot by the original pLDDT color gradation.

Although the number of proteins in PDB and NA groups are similar to each other, the number of variations within these groups notably differ. The variant count in the PDB group was nearly double of that in the NA group (Fig. 1b), implying a bias of ClinVar towards reporting more instances of proteins with known structures (PDB group) compared to proteins without any structures (NA group). The entire dataset having a distribution of 40%-60% pathogenic-benign labels is relatively balanced. However, the distribution of the pathogenicity classes vary across the PDB and NA groups (Fig. 1b). As such, the NA group harbors mostly neutral mutations and much fewer pathogenic variants than does the PDB group (Fig. 1b), also signifying a bias of ClinVar towards reporting more pathogenic variants from proteins with known structures.

To address whether or not there are any distinctions in these protein sets possibly sourcing these biases, we compared the length and the level of functional annotations of the proteins in PDB and NA groups (Fig. 1c). While length distributions were highly similar to each other, the number of annotations were not. Specifically, the proteins in the PDB group have higher number of annotations than the proteins in the NA group (Fig. 1c). The large right-skew in the PDB group is likely a result of the well-recognized bias of biomedical databases towards richly annotated genes [23, 39]. Many databases, including PDB, have a tendency to report more instances of richly annotated genes compared to the ones with relatively fewer annotations [23, 39–41]. Hence, the large difference in the missense variant counts and pathogenicity distributions between PDB and NA groups could be due to a bias of ClinVar, i.e. its submitters, toward reporting variations from proteins with high functional annotations and/or known structures.

We note here that, even if the associated PDB structure is partial and the variant in question is not structurally characterized, it was still counted in the PDB group in the Fig. 1a. Often experimental structures, particularly from crystallization experiments, contain unmodeled regions with sizes ranging from a few amino acids to large domains [42]. Thus, being associated with any PDB entry does not necessarily suggest the full-length structural characterization of a given protein. Herein, Titin, one of the longest proteins in the human proteome, can be visited as an example. Despite its extensive structural characterization resulting in many distinct PDB entries, to date only a small portion of its structure, less than 2%, could be resolved. This is significant because if we were to count the structurally characterized and uncharacterized variants, their distributions would likely appear less distinct from each other than shown in Fig. 1b. Regardless, this analysis underlined that a significant portion of human variations cannot be analyzed by a structure-based predictor due to their missing/incomplete structural characterization.

### Reviewing the Experimental and AF2 Computed Protein Structures of the Dataset

AF2 computations provide a confidence score for each residue, pLDDT, which is based on the IDDT metric allowing one to assess the reliability of local structures [43]. We delineated the statistics of pLDDT distributions of the AF2 structures in this dataset (Fig. 1d). Only, a subtle increase in the central tendency measures were noted for the PDB group compared to the NA group, underscoring a slightly more favorable AF2 prediction for the proteins with known structures. Other than this difference, pLDDT distribution statistics were high similar to each other for the proteins in the PDB and NA groups, indicating indifference of AF2 confidence to PDB availability. However, we reported that the median and mean measures were notably higher in the soluble proteins and enzymes (Fig. S2).

A skewness-kurtosis plot was further used to examine the shape of each pLDDT distribution from the PDB and NA groups (Fig. 1e). This analysis suggested that most of the structures in this dataset have a negatively skewed and light-tailed (leptokurtic) pLDDT distributions. Two example pLDDT distributions and their corresponding AF2 structures were also depicted (Fig. 1e). These plots compare two pLDDT distributions: green with extreme skewness/kurtosis and blue with normal-like features, both marked on a separate skewness-kurtosis plot (Fig. 1e). The corresponding AF2 structures of these distinct pLDDT distributions were also visualized on the right side of the plots.

The AF2 predictions of this dataset, covering the full-length structures, overall exhibit a low frequency of low confidence regions, highlighting the impartiality of AF2 predictions to the availability of an experimental structure for this dataset. Overall, AF2 structures offer the full-length representation of a protein along with a confidence score for each amino acid. Thus, the use of AF2 structures may provide an advantage to structure-based pathogenicity predictors, relieving their PDB dependency to the variant/protein in question and broadening their coverage.

### Construction of the AFFIPred Classifiers

We split 7,368 proteins into three equal groups, ensuring similar characteristics such as length, annotations, protein types within each group (Fig. S3a). Next, we created three distinct sets of protein variations from these splits, ensuring no protein appeared in multiple sets (Fig. S3b). These variations were also comparable in terms of pathogenicity and review status. We trained AFFIPred models on two-thirds of the ClinVar variations (∼50,000) using a nested cross-validation (CV) approach (Table 1). Nested CV approach is preferred over CV or single validation approaches as it provides a more robust estimate of model performance [44]. However, even with nested CV, proteins present in both training and validation sets can bias the results. For instance, inclusion of the proteins that harbor only one class of variants, i.e. pathogenic or neutral both in the training and validation sets can lead to overestimated performance [45, 46]. To address this potential bias, we evaluated the final models on the unseen test sets generated previously (Fig. S3b). This approach eliminates protein-based bias and allows for an unbiased assessment of AFFIPred’s performance compared to other classifiers. In line with this expectation, we observed a decrease in performance of AFFIPred on the unseen test sets compared to that in the nested CV, suggesting the potential impact of common proteins in the training and validations datasets of nested CV (Table 1).

**Table 1.** AFFIPred’s Performance in nested CV and against unseen test set without common proteins in the training set.

|  | Nested CV |  | unseen |  |
| --- | --- | --- | --- | --- |
|  | size | AUC | size | AUC |
| AFFIPred1 | 49301 | 0.953±0.001 | 26029 | 0.921 |
| AFFIPred2 | 49421 | 0.954±0.001 | 25909 | 0.916 |
| AFFIPred3 | 51938 | 0.956±0.001 | 23392 | 0.915 |

SHAP values were used to assess the importance of each feature used by the AFFIPred classifiers [47]. This analysis reflected the importance of not only the well-known sequence-based features such as PSIC scores [3, 48] but also all three AF2-based structure attributes including the solvent accessible surface area (SASA) and pLDDT scores of the native positions (Fig. S3c). Among these, SASA is a well-established structural feature linked to pathogenicity [7, 8, 10], while pLDDT was newly identified as relevant to pathogenicity [7, 49]. Notably, pathogenic variants tended to have lower SASA and higher pLDDT values (Fig. 1b-c). We also reported that average pLDDT of the selected AF2 structure was also an important feature for all three classifiers. Average pLDDT was slightly higher for the pathogenic cases compared to neutral variants. Distribution of the feature values across pathogenicity labels were given in Fig. S4.

### Performance Assessment on Unseen Test Sets

We compared the AUC-based performance of AFFIPred against 40 classifiers/scores (Figs. 2a and S5). AFFIPred showed a comparable or better performance than the non-integrated tools, reaching an AUC of 0.92 on an unseen test set that was formed by distinct proteins from the training dataset. Two scores, AlphaMissense and VEST4 surpassed AFFIPred with AUC values close to 0.95 (Fig. 2a). Most of the integrated predictors, like BayesDel, ClinPred, and MetaRNN, achieved near-perfect AUC scores. We note that the training data of the tools other than AFFIPred and Rhapsody were not inspected to eliminate any possible overlaps with the test set, reflecting the plausibility of overestimated AUC for the methods other than AFFIPred and Rhapsody. Performance of Rhapsody was reported for the variants that were not found in its training data, while the performance of AFFIPred was further documented for the variants and proteins that were not found in its training dataset. Reversely, ClinPred that was trained on variants collected from the ClinVar [50], likely exhibited an inflated performance due to the possible overlap between its training data and our test set.

**Fig 2.**
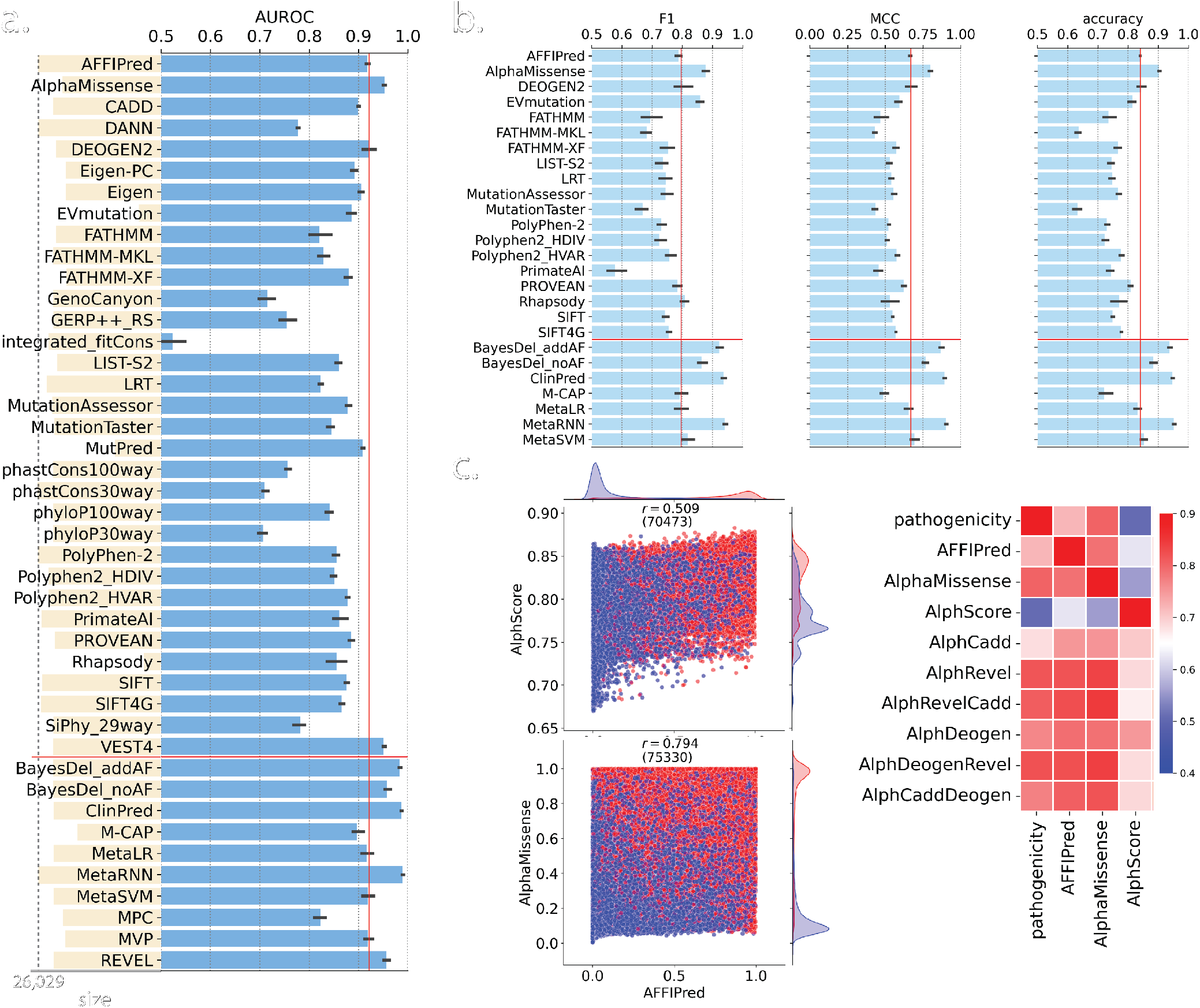
Performance Assessment. Classification performance based on (a) AUROC, (b) F1, MCC and accuracy was plotted. Panel (a) also shows the number of variations that were predicted by each tool by orange bars. Dotted line marks the predicted variation size for AFFIPred. Red vertical line marks the performance of AFFIPred. Integrated scores were shown at the bottom separated by a horizontal red line. predictors were shown on the right. (c) Joint plots show the distribution of the AFFIPred scores with respect to AlphScore [7] and AlphaMissense [24] scores, which employ AF2 structures and/or use AF2-derived features for pathogenicity prediction. Pathogenic variants were marked by red circles and benign variants were marked by blue circles. (d) Heatmap shows Spearman rank correlation coefficient between pathogenicity and pathogenicity scores from AFFIPred, AlphaMissense, AlphScore and integrated versions of AlphScore.

We further compared the AFFIPred’s performance against the tools that predicted a classification outcome using F1 score, Matthew’s correlation coefficient (MCC), and accuracy metrics (Fig. 2b). AFFIPred consistently ranked in the top three methods among non-integrated predictors for all these metrics (Fig. 2a-b). Notably, AlphaMissense was the most powerful method that also utilizes AF2 structures [24] in all four metrics [51, 52]. Figs. 2a and S6 also show the coverage of each predictor, i.e. the number of predicted variants. AFFIPred and PolyPhen-2 predicted all variants. While all other methods missed some, Rhapsody predicted around a third of the dataset, exemplifying the limitation of structure-based classifier due to reliance on the availability of an experimental structure.

We further evaluated AFFIPred’s performance against new scores called AlphScore, which relies solely on the AF2 structural features [7] and AlphaMissense which uses the AlphaFold’s system and trained on a large variant dataset [24]. Particularly, these comparisons suggested a medium level correlation between AlphScore and AFFIPred while a strong one between AlphaMissense and AFFIPred. Score distributions further revealed a similarity between AFFIPred and AlphaMissense predictions while a distinction in AFFIPred and AlphScore (Figs. 2b). As such, score distributions across pathogenicity classes implied that AFFIPred and AlphaMissense were able to distinguish pathogenic and benign variants more effectively than did AlphScore (Fig. 2c). Additionally, Fig. 2d demonstrates a strong correlation between AFFIPred scores and the integrated scores of AlphScore with Revel, CADD or DEOGEN, which themselves are highly correlated with pathogenicity (Fig. 2d). In contrast, AlphScore exhibited weaker correlation with both pathogenicity and the integrated scores. In summary, AFFIPred classifiers, combining sequence and AF2 structural information, achieve a comparable, if not superior, pathogenicity prediction performance and are not restricted by PDB availability.

### Comparison of Structural Features from Experimental and AF2 Structures

We conducted a detailed comparison of the structural features extracted from PDB and AF2 structures to assess their consistency (Fig. 3). Specifically, the pLDDT and rASA distributions that were collected from the AF2 structures were examined across pathogenic and benign variants (Fig. 3a). This analysis revealed a unimodal distribution of both scores in the pathogenic variants, while the distributions of the neutral variations had bimodal shapes. These observations were true for the entire dataset (ClinVar), the NA and PDB group of variants (Fig. 3a). However, both SASA and pLDDT scores in the variant set that could be predicted by Rhapsody converged to a unimodal distribution, indicating the loss of the lower/upper tail peaks in the neutral group. Since Rhapsody requires a corresponding PDB structure for prediction, the variants it misses are likely not located in the structurally characterized regions. These missing variants tend to be neutral with low pLDDT and high SASA scores, readily implying a bias in the training dataset compiled solely relying on structurally determined variations. Furthermore, the discriminatory power of these structural features, pLDDT and SASA, which are among important features for our (Fig. S3) and many other models [7, 8, 49], appears to be diminished for the variations with structural characterization (Fig. 3a). We also checked the WT PSIC score in the same variant sets, and observed that the eliminated group of variants in the Rhapsody predicted set did not have a particular pattern in their PSIC scores, as they did in their SASA and pLDDT scores (Fig. 3a).

**Fig 3.**
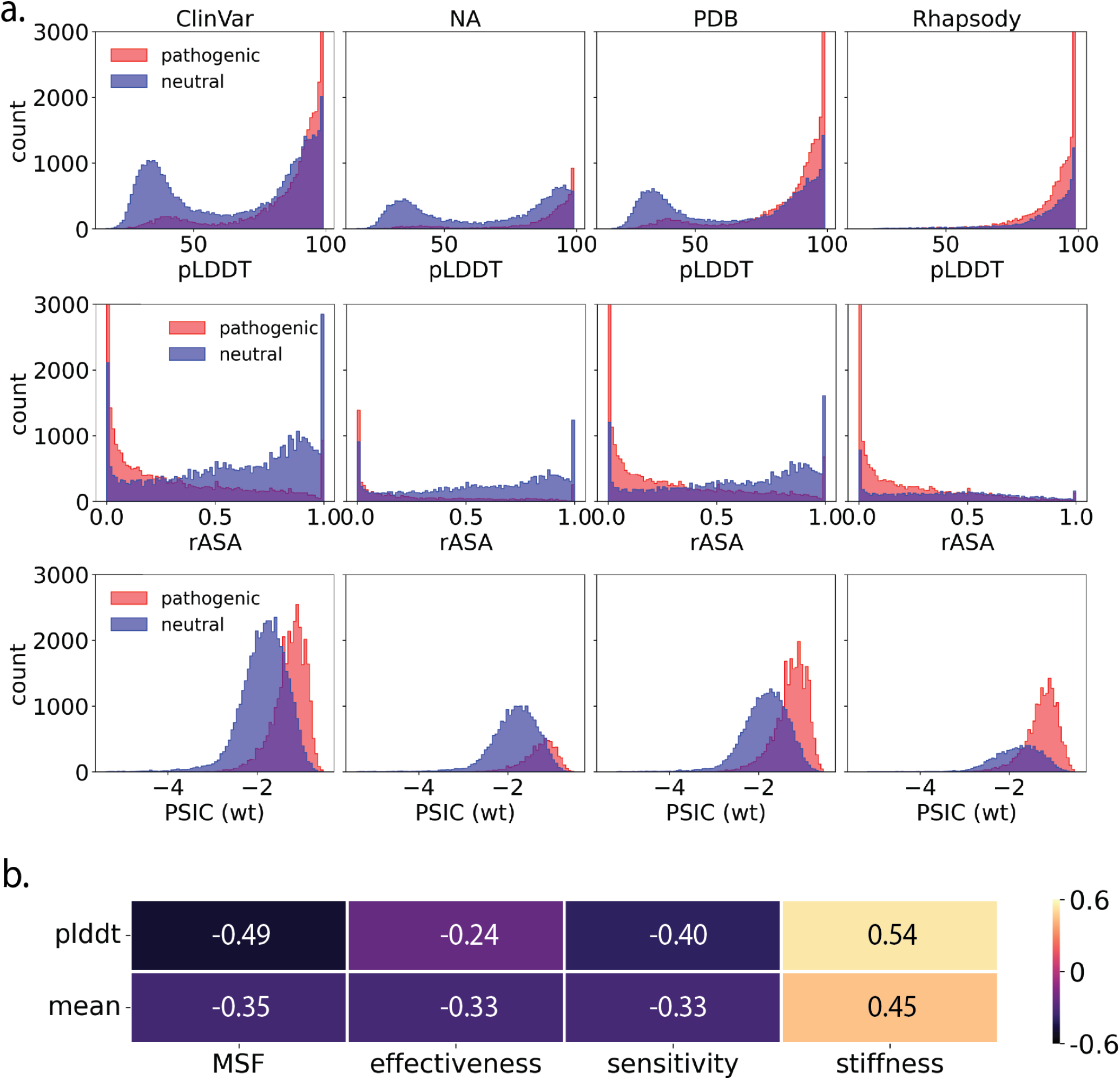
Comparison of Structural Features. (a) Distribution of pLDDT, relative SASA and WT PSIC scores of the variations in the entire dataset (ClinVar), in the NA, PDB groups and in the Rhapsody predicted set. (b) Pearson correlation coefficient between pLDDT scores of variants and mean pLDDT and dynamic features of Rhapsody that were calculated from PDB structures.

We investigated the structural attributes of Rhapsody, extracted from dynamic structures through coarse-grained molecular dynamics simulations [6, 8]. These attributes primarily include mean square fluctuations (MSF), effectiveness, sensitivity, and stiffness (Fig. 3c). Upon examining the relation of the dynamic features with the pLDDT-based features of our models, a low-level correlation was spotted, implying a partial explanation of the dynamic features by pLDDT-based features. Hence, we reported that the pLDDT scores remain valuable for understanding the solvent accessibility and dynamics of experimental structure, albeit to some extent.

We also plotted absolute and relative SASA values for a set of variants that were structurally characterized and thus predictable by Rhapsody. Essentially, Rhapsody computes absolute SASA values, while we relied on relative SASA, representing the ratio of absolute SASA to the maximum SASA of the amino acid. Both absolute and relative SASA measurements are interpreted similarly as such higher values denote solvent-exposed regions, while lower values indicate buried regions [33]. Despite their shared interpretation, we did not observe a strong association between the SASA and rASA calculations from PDB and AF2 structures, respectively (Fig. 4a). Strikingly, there were a number of inconsistencies between SASA and rASA measurements that some data points were spotted scattered in the upper-left and bottom-right corners of Fig. 4a, indicating low absolute SASA but high relative SASA (or vice versa).

**Fig 4.**
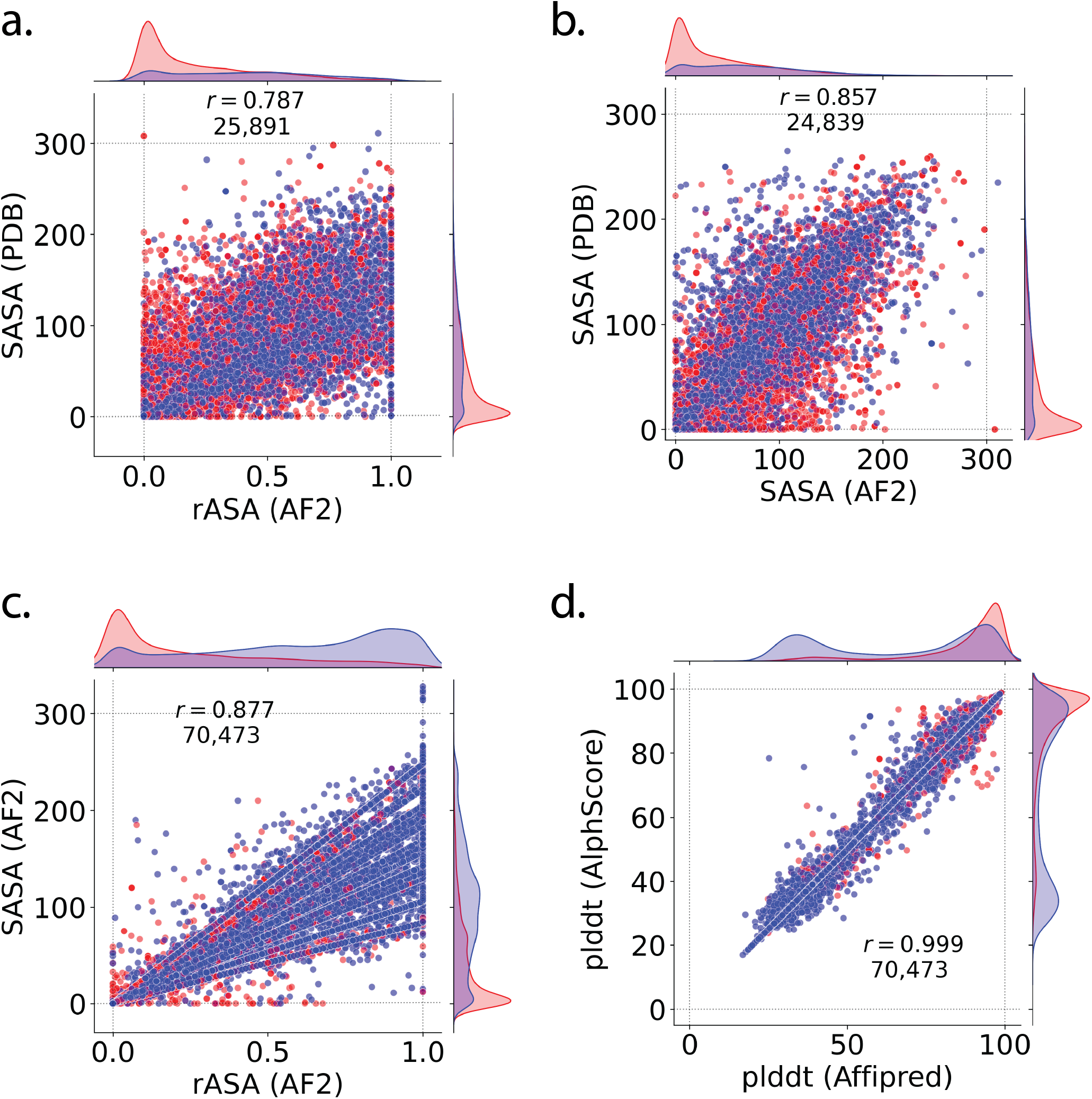
Absolute versus Relative SASA. (a) Comparison of relative and absolute SASA values that were calculated from AF2 structures by AFFIPred and PDB structures by Rhapsody, respectively. (b) Comparison of absolute SASA values that were calculated from AF2 structures by AlphScore [7] and PDB structures by Rhapsody, respectively. (c) Comparison of relative and absolute SASA values that were calculated from AF2 structures by AFFIPred and AlphScore. (d) Comparison of pLDDT scores calculated by AFFIPred and AlphScore. Pearson correlation coefficients and total number of data points were given for each comparison.

We also compared the absolute SASA computations by Rhapsody from PDB structures with those derived from AF2 structures by the AlphScore method [7] (Fig. 4b). This comparison for ∼24,000 variants showed a stronger correlation given the absence of SASA scaling (Fig. 4a). However, the expected close association was still absent and some inconsistencies were still present (Fig. 4b). Comparison of the absolute and relative SASA values from the AF2 structures showed an even stronger association for more than ∼70,000 variants (Fig. 4c). In fact, the scaling of absolute SASA to the maximum possible values can be distinguished in the Fig. 4c as straight lines for each amino acid. Despite an increase in the strength of the association, some inconsistencies were still present at the bottom-right of Fig. 4c. As these points with low absolute SASA and high relative SASA were delineated, we found out that they were mostly the variations from the C-termini of particularly long proteins such as DNAH10. Such long proteins have overlapping AF2 structures and thus selection of the AF2 structure for feature extraction could have led to these inconsistent SASA measurements.

The differences in the absolute and relative SASA measurements could stem from two factors. One is the scaling of SASA values to their maximum possible values. The other, and more significant factor based on the correlation strength and the number of variants analyzed in Fig. 4a-c, is the inherent structural discrepancies between the AF2 and the experimental structures. Essentially, this finding raises the question of whether AF2 structures can complement experimental structures for calculating structural attributes, warranting further investigation for the discrepant points.

We lastly examined the pLDDT scores obtained here and from AlphScore calculations by Schmidt et al. [7] (Fig. 4d). This comparison exhibited an almost perfect correlation, underscoring the agreement of pLDDT extraction by the AFFIPred and AlpScore. Fig. 4d shows that some data points deviate from the expected perfect correlation. These deviations might be caused by a few factors such as version difference in AF2 structures. Our calculations used AlphaFold2 v4 models, while the AlphScore calculations might have used a different version. Second is the selection of AF2 structures for the proteins exceeding 2700 amino acids may have introduced some discrepancies in the pLDDT scores.

### Challenges Associated with the Usage of PDB Structures for Feature Extraction

Our analysis highlighted inconsistencies between AF2 and PDB structures, particularly for SASA computations. To address whether or not, these inconsistencies are due to any miscalculation on the Rhapsody’s and/or our side, we selected six variants from four proteins for which absolute SASA calculations by Rhapsody using PDB structures and relative SASA calculations by AFFIPred using AF2 structures significantly differed.

The first inconsistency involves the variants of Y145 from Syntaxin-binding protein 1 (Fig. 5a). The AF2 structure for Munc18-1 has a high accuracy (mean pLDDT of 89.8) and low predicted aligned error (PAE) (Fig. 5a), implying a reliable prediction. Accordingly, AF2 predicts Y145 to be buried within the protein with a zero rASA (Fig. 5a). In turn, Rhapsody, which utilized the only PDB structure (6L03) that captured a tiny fragment (8 amino acids) of Munc18-1 (595 amino acids) [53], calculated the absolute SASA of Y145 as 308 ^°^*A*^2^, which is even larger than the maximum SASA of a tyrosine (263) [33] (Table 2). The Munc18-1 fragment, which forms a complex with a tyrosine phosphatase, is phosphorylated at the Y145 position in the PDB complex (Fig. 5a). Here it is worth noting that analyzing the entire PDB complex might in part counteract the overestimation of absolute SASA of Y145, because the phosphorylated Y145 in the small Munc18-1 fragment is completely buried in the complex (Fig. 5a). However, Rhapsody recruits single chains from complex structures, a feature which was deemed to be improved in future models [6].

**Table 2.** Prediction results of 6 cases with varying SASA computations.

| gene | variant | rASA<sup>†</sup> | SASA<sup>‡</sup> | Class<sup>§</sup> | Rhapsody | Affipred |
| --- | --- | --- | --- | --- | --- | --- |
| STXBP1 | Y145C | 0.0 | 308(263) | P** | neutral | pathogenic |
|  | Y145H |  |  | P* | neutral | pathogenic |
| TP53 | D49N | 1.0 | 1(193) | B*** | neutral | benign |
|  | D49H |  |  | B*** | deleterious | benign |
| AKT3 | W79C | 1.0 | 25(285) | P* | neutral | benign |
| PRDM16 | L151V | 1.0 | 34(201) | B* | prob.delet. | benign |

<sup>†</sup> Relative SASA from AF2 structures <sup>‡</sup> Absolute SASA from PDB structures, max.SASA [33] is given in parenthesis. <sup>§</sup> P: pathogenic, B: benign. \* indicates review stars of the variants

**Fig 5.**
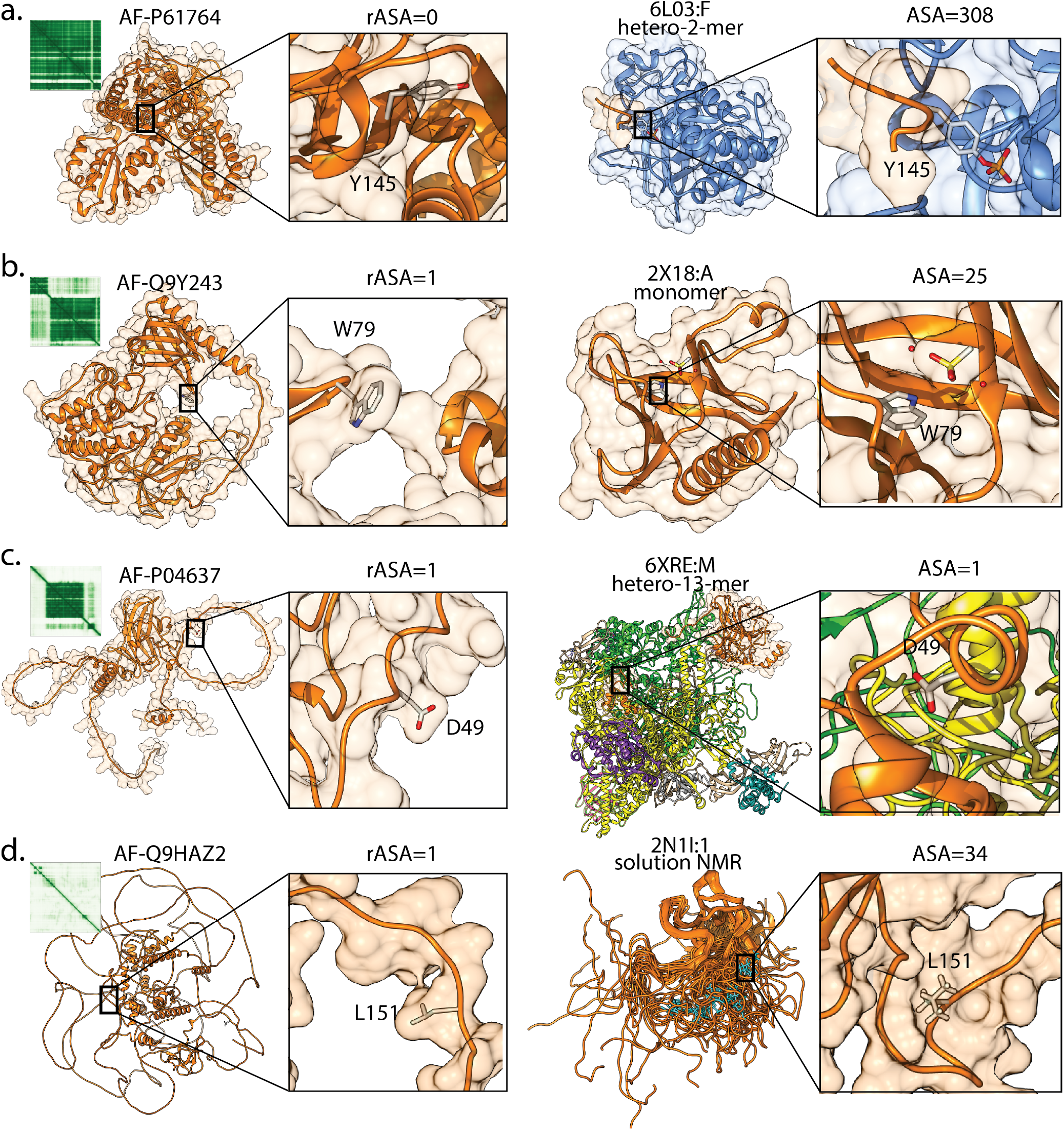
AF2 versus PDB structures. Four protein cases of (a) Syntaxin-binding protein 1, (b) RAC-gamma serine/threonine-protein kinase encoded by AKT3 gene, (c) Cellular tumor antigen p53 and (d) PRDM16 were structurally visualized. The selected position was rendered in licorice and shown on the global structure which is illustrated as orange cartoon and surface renders. AF2 structure of the given position were shown on the left and the PDB structure that were employed for Rhapsody predictions were shown on the right. A close-up view is shown to visualize the accessible surface area of the positions for both structures, labeling relative SASA for AF2 structures, and absolute SASA for PDB structures. PAE plots were also shown for AF2 structures.

The second example to the inconsistent SASA measurements is the position W79 of RAC-gamma serine/threonine-protein kinase. The AF2 structure for this 480-amino-acid protein has a high predicted accuracy (mean pLDDT score of 81.8) with low error throughout most of the sequence, except for a central loop (Fig. 5b). However, the available PDB structure only captures the N-terminal domain (100 amino acids). In this case, both PDB and AF2 structures show W79 to be fully exposed. However, the crystal structure (2X18) contains a sulfate molecule near the W79 sidechain, affecting the SASA calculation from the PDB structure (Fig. 5b). This highlights that the presence of small molecules such as ligands and buffer contaminants in the PDB structure can influence SASA calculations.

The third case involves the cellular tumor antigen p53. We examined a variant position (D49) within its trans-activation domain (TAD) (Fig. 5c). TADs are typically disordered but can adopt transient ordered structures upon binding to other proteins [54, 55]. The AF2 structure of p53 predicted D49 to be fully exposed, reflecting its intrinsic disorder. However, the available PDB structure representing p53 bound to RNA polymerase II showed D49 as buried. This discrepancy arises because the PDB structure captures a complex wherein its TAD adopts a transient secondary structure burying the D49 sidechain. While the AF2 structure represents the free protein. In one aspect, SASA calculation from the multimeric complexes would be meaningful for understanding their functional impact [56, 57]. However, regarding these cases, we noted that the complex PDB structures may or may not misrepresent the solvent accessibility of the free form of the variant. Hence, direct recruitment of PDB structures for SASA calculations would impede with the systematical analysis of the free structures. In this case, the AF2 structure offered an advantage by providing the full-length, unbound structure for p53.

The fourth case examines the variant position L151 in PRDM16, a protein involved in fat cell development (Fig. 5d). The AF2 structure predicts L151 to be fully exposed. However, the available NMR structure shows L151 as less exposed. This difference arises because NMR structures capture multiple snapshots of a protein’s flexible regions. In this case, L151 resides in a flexible loop, and its exposure varies among twenty solution models. Thus, selecting a single NMR snapshot might not accurately reflect its true accessibility. AF2 structures, on the other hand, provide a single, static structure with a better representation of the NMR ensemble, particularly IDRs. While the mean pLDDT scores and PAE plot reflected that the AF2 predicted structure of PRDM16 is far from perfect, it still provided a more realistic representation of L151’s solvent accessibility.

We examined six protein variants from four positions using both PDB structures and AF2 structures in Table 2. All of these variations represent a problematic case wherein PDB structures failed to represent the SASA of the native position. Rhapsody, a state-of-the-art structure-based pathogenicity classifier, wrongly predicted 5 of 6 cases, implying a dramatic reduction in its performance compared to Fig. 2a-b. On the other hand, AFFIPred correctly predicted five of these six variations and misclassified only W79C (Table 2, leading to a performance that aligns well-with its general performance (Fig. 2a-b). We stress that the reduction of Rhapsody’s performance in these six cases is likely to be related to the problems in the corresponding PDB structures but not in the tool itself [6, 8]. Exemplifying through the Rhapsody’s PDB selection approach, we reported that the structure-based classifiers that rely on PDB structures could misinterpret the structural characteristics of a given variant position due to (i) recruitment of a single chain from a multimer, (ii) post translational or chemical modifications, (iii) presence of non-proteinous molecules, and/or (iv) missing regions close to the variant position of interest. While other conditions may also lead to misinterpretation of structural features of variants, these explained e most of the inconsistent SASA calculations, spotted in the Fig. 5.

### AFFIPred Classification for All Variations of Human Proteome

94% of human proteome enumerating a total of 19,365 proteins and corresponding 20,400 AF2 structures (Fig. 6a) were processed to generate a total of 210,156,568 variations. The reason for the missing portion (6%) was due to lacking of precomputed PSIC profiles. All these variations were predicted by three AFFIPred classifiers. Overall, around one quartile of the variations were found pathogenic while the remaining variations were labeled as neutral. Given three distinct predictions by three AFFIPred classifiers, we also devised a confidence rank based on the average and standard deviation of the AFFIPred probability values (Fig. 6b). The variations with a mean probability value between [0-0.1] or [0.9-1.0] were assigned with three stars; the variations with a mean probability score (0.1-0.4) or (0.6-0.9) and a standard deviation less than 0.2 were assigned with two stars; and the rest of mutations were given a single star (Fig. 6b). Distribution of the confidence stars were given in the Fig. 6b. Nearly half (48%) of the predictions received a three-star rating, suggesting strong agreement among three AFFIPred models on the variant’s pathogenicity. Conversely, ∼13% of the cases were assigned a one-star rating, indicating high variability in their pathogenicity prediction.

**Fig 6.**
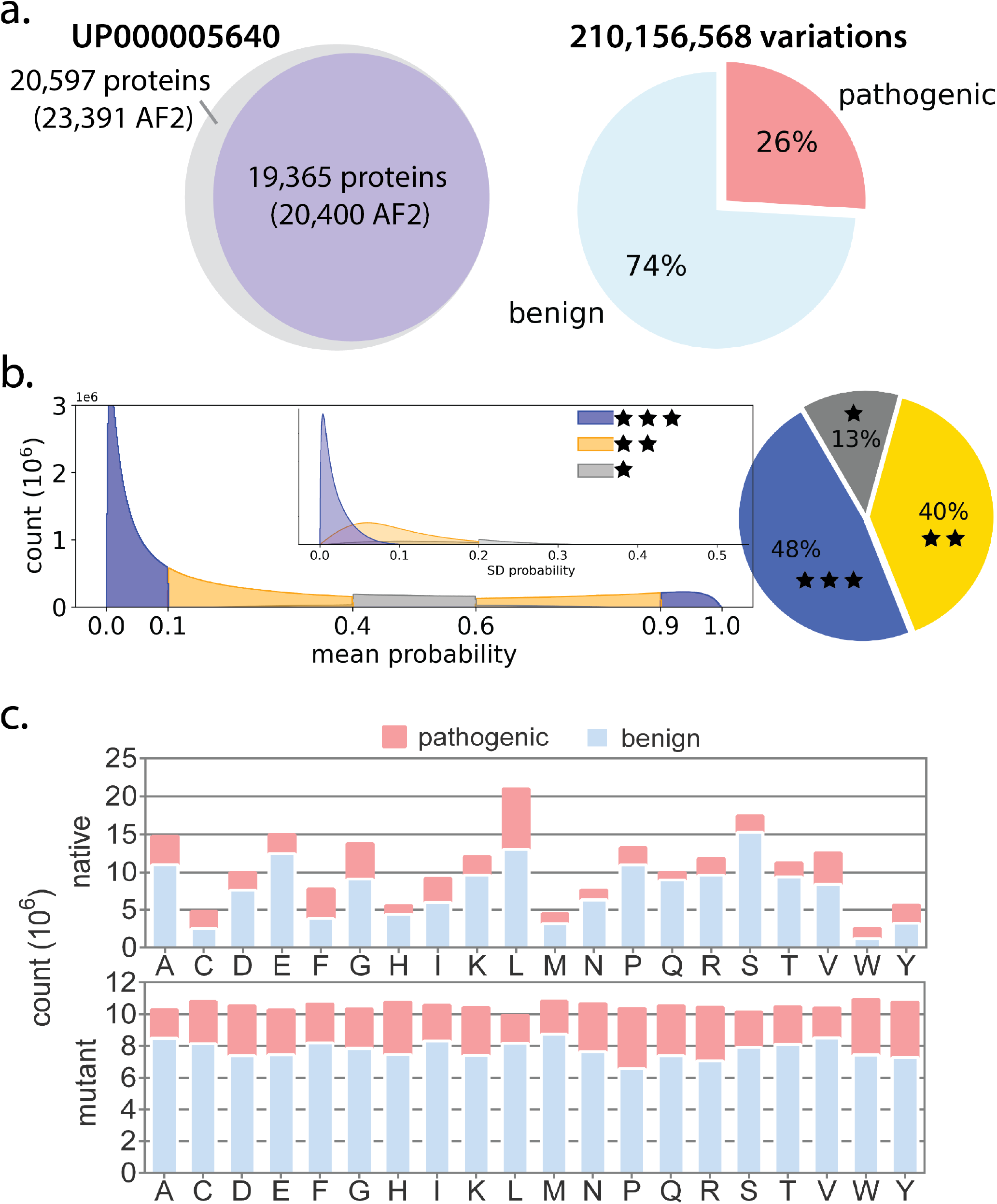
AFFIPred Prediction of the Human Proteome. (a) Almost all proteins in the human proteome enumerating a total of 19,365 distinct Uniprot identifiers were analyzed by processing 20,400 AF2 structures. Pathogenicity of over 210 million human variations were predicted by three AFFIPred classifiers. (b) Distribution of the confidence ranks that were assigned to each variation based on the mean and standard deviation (inset) of the AFFIPred probability were given. (b) Distribution of pathogenicity classes across amino acids in the native and mutant positions of the human proteome.

All possible variations in the human proteome showed a highly unbalanced distribution of amino acid frequencies in the native positions (Fig. 6c), an observation which is in line with the amino acid composition of human proteome [58]. Aromatic amino acids (W, F, Y) and C were the least tolerant amino acids while polar amino acids like N, Q, S and T had the least pathogenic mutation fraction, implying their tolerance for variations. We also plotted the neutral and pathogenic mutation distribution in the mutant positions. Particiarly W, Y, P and R were the most deleterious amino acids, causing a pathogenic label in the human proteome. On the other hand, substitutions of small aliphatic amino acids of A, I, L and V were mostly benign. These predictions can be accessed through the web or command line interface of AFFIPred (https://github.com/timucinlab/AFFIPred), that accepts queries based on inputs of protein, genomic locations, variant identifier or vcf files.

## Conclusion

Traditionally, predicting pathogenicity has relied heavily on analyzing protein sequences, likely because obtaining a protein sequence is significantly cheaper and easier than determining its 3D structure. However, this preference for sequence-based methods might not reflect an inherent superiority of sequence over structure for pathogenicity prediction. Overall, dependence on the PDB characterized variants would lead to particular exclusion of the set of neutral variants with low pLDDT and high SASA, source a possible bias. Furthermore, the structural features such as pLDDT and SASA may not be as informative as they would be in an AF2-based classifier.

Notwithstanding the power of structural information, structure-based predictors have certain limitations. They rely on the knowledge obtained from the variations with known structures, missing those in the IDRs. Further, they depend on experimental structures which may have certain shortcomings, leading to misinterpretation of the structural characteristics of a given variant. As exemplified here by a number of cases, some conditions might result in a wrongful calculation of SASA of a given variant, reflecting the need for additional pre-processing steps for direct recruitment of a PDB structures for extracting of structural information.

This study introduces a novel structure-based pathogenicity classifier, AFFIPred, that merges both sequence and AF2-based structural information. This combination tackles limitations of traditional structure-based classifiers while maintaining accuracy. AFFIPred implementation that utilizes AF2 predictions that are currently accessible and provide full-length representations ofproteins in their unliganded state allowed more precise SASA computations than those based on PDB structures. We surmise that AFFIPred is a novel structure-based predictor, paving the way for the development of a wider range of structure-based predictors and for the transformation of traditional PDB-dependent methods.

## Supporting information

### S1 Hyperparameter Optimization

Optimized parameters and their search space was shown. The optimized value for each parameter was given at the top of each plot. Panels illustrate the optimization results for three training procedures.

### S2 Distribution of pLDDT Statistics

(a) Soluble and membrane proteins, (b) proteins with and without an EC number

### S3 Dataset Construction and Feature Importance

(a) Proteins forming the ClinVar dataset was randomly divided into three distinct groups. Distribution of protein length, number of GO annotations, and average plddt scores were visualized for the entire set and each protein subset. Heatmaps illustrate the relative proportions of enzymes, membrane proteins, and proteins with PDB structures in each protein subset. (b) After collection of the variations from each protein subset, three distinct variation datasets were formed. Heatmaps displayed the relative proportions of the review stars and pathogenicity labels. (c) Feature importance of each classifier was assessed by SHAP values. Red indicates high, blue indicates low values of features. Model number reflects the unseen test set that was used to assess the performance. AFFIPred1 was trained on the combination of second and third variation sets and tested on the first variation set.

### S4 Distribution of the Feature Values Across Pathogenicity Labels

### S5 AUC Results of the Benchmarking against 40 Pathogenicity Predictors

Converted rank scores were used to plot ROC curves. Three panels show benchmarking of 40 tools including AFFIPred on three distinct test sets that was unseen for the AFFIPred.

### S6 Performance versus Coverage

Plots show performance; AUC, F1, MCC or accuracy on the y-axis and the number of predicted variations on the x-axis. Three columns represent the assessments based on three distinct test sets, which was unseen for the AFFIPred and do not have any overlapping proteins.

