## Supplementary material for "AFFIPred: AlphaFold2 Structure-based Functional Impact Prediction of Missense Variations": SI file

### List of Figures

1 **Hyperparameter Optimization.** Optimized parameters and their search space was plotted. The optimized value for each parameter was given at the top of each plot. Rows illustrate the optimization results from three training procedures. . . . . . . . . . . . . . . . 2

2 **Distribution of pLDDT Statistics.** (a) Soluble and membrane proteins, (b) proteins with and without an EC number . . . . . . . . . . . . . . . . . . . . . . . . . . . . . . . . . . . 3

3 **Dataset Construction and Feature Importance** (a) Proteins forming the ClinVar dataset was randomly divided into three distinct groups. Distribution of protein length, number of GO annotations, and average plddt scores were visualized for the entire set and each protein subset. Heatmaps illustrate the relative proportions of enzymes, membrane proteins, and proteins with PDB structures in each protein subset. (b) After collection of the variations from each protein subset, three distinct variation datasets were formed. Heatmaps displayed the relative proportions of the review stars and pathogenicity labels. (c) Feature importance of each classifier was assessed by SHAP values. Red indicates high, blue indicates low values of features. Model number reflects the unseen test set that was used to assess the performance. AFFIPred1 was trained on the combination of second and third variation sets and tested on the first variation set. . . . . . . . . . . . . . . . . . . . . 4

4 **Distribution of the Feature Values Across Pathogenicity Labels.** . . . . . . . . . . 5

5 **AUC Results of the Benchmarking against 40 Pathogenicity Predictors.** Rank scores were used to plot ROC curves. Three panels show benchmarking of 40 tools including AFFIPred on three distinct test sets that was unseen for the AFFIPred. . . . . . . . . . . 6

6 **Performance versus Coverage.** Plots show performance; AUC, F1, MCC or accuracy on the y-axis and the number of predicted variations on the x-axis. Three columns represent the assessments based on three distinct test sets, which was unseen for the AFFIPred and do not have any overlapping proteins. . . . . . . . . . . . . . . . . . . . . . . . . . . . . 7

### Supplementary Figures

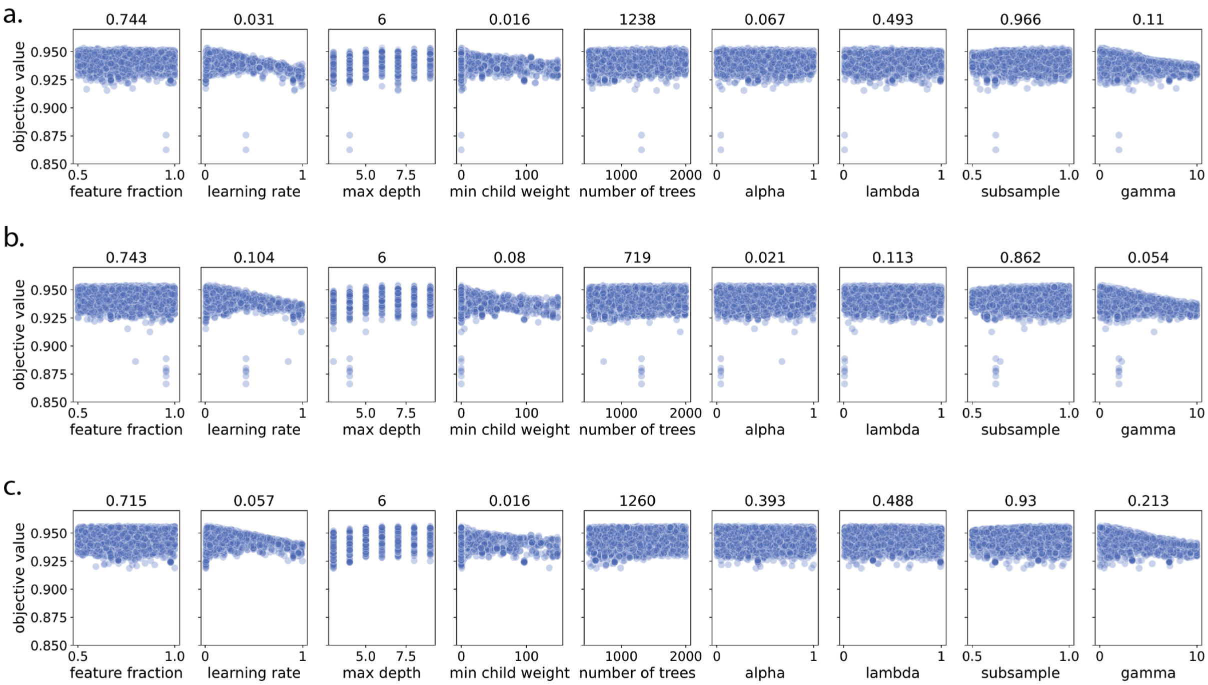


**Fig 1. Hyperparameter Optimization.** Optimized parameters and their search space was plotted. The optimized value for each parameter was given at the top of each plot. Rows illustrate the optimization results from three training procedures.

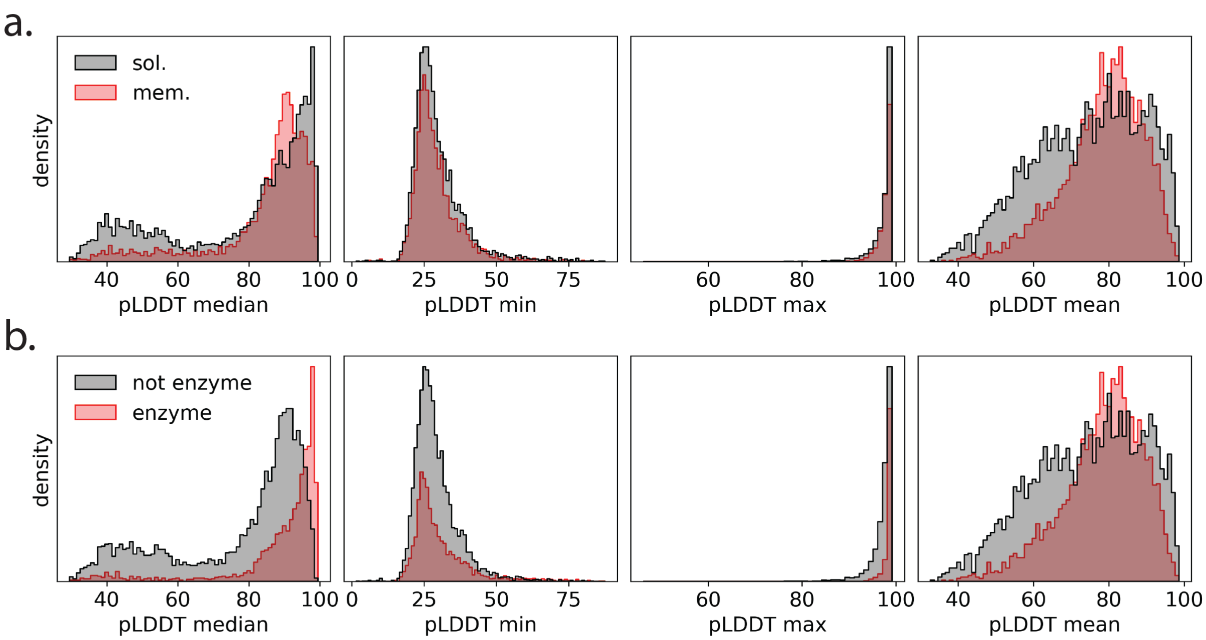


**Fig 2. Distribution of pLDDT Statistics.** (a) Soluble and membrane proteins, (b) proteins with and without an EC number

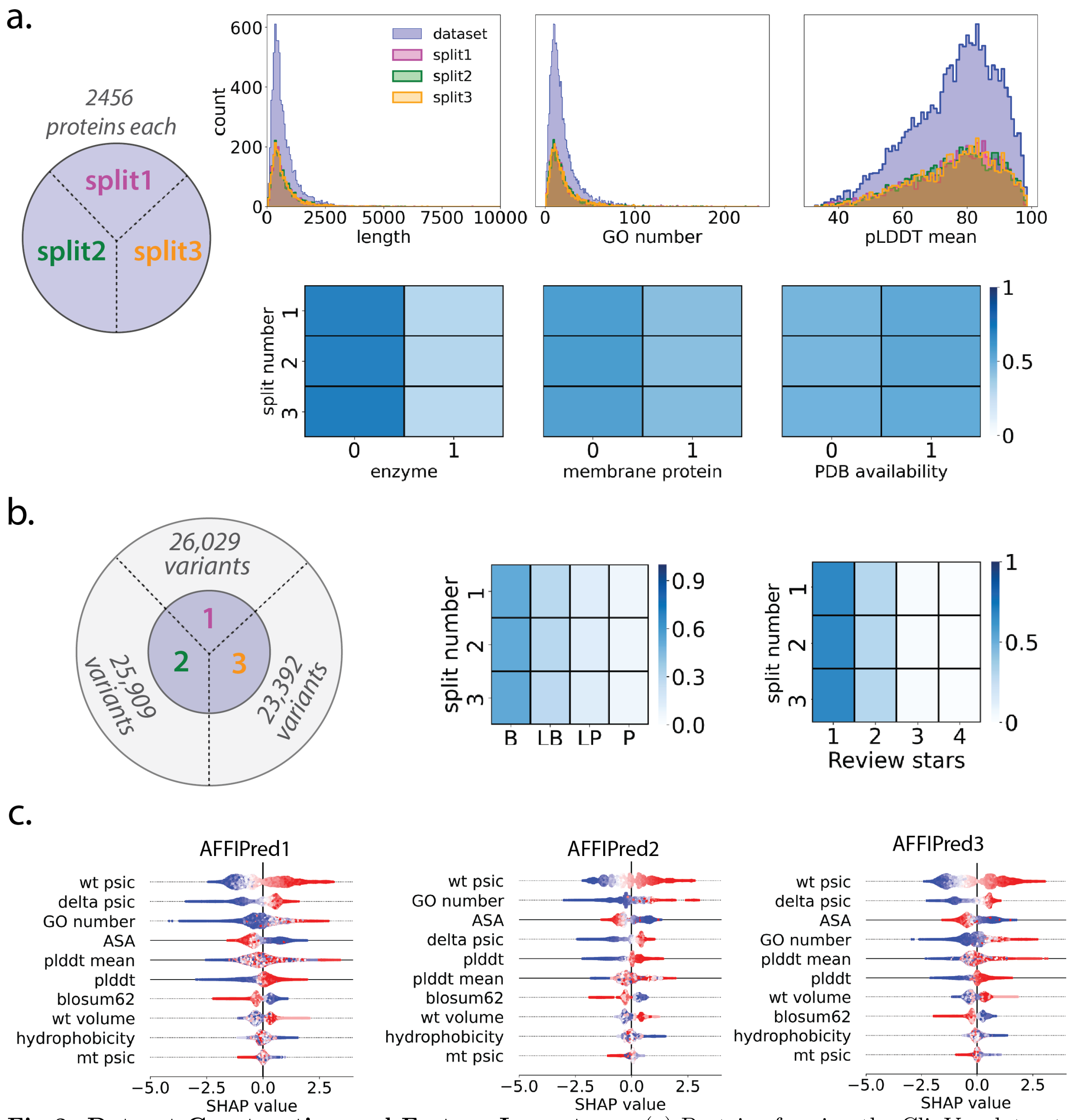


**Fig 3. Dataset Construction and Feature Importance** (a) Proteins forming the ClinVar dataset was randomly divided into three distinct groups. Distribution of protein length, number of GO annotations, and average plddt scores were visualized for the entire set and each protein subset. Heatmaps illustrate the relative proportions of enzymes, membrane proteins, and proteins with PDB structures in each protein subset. (b) After collection of the variations from each protein subset, three distinct variation datasets were formed. Heatmaps displayed the relative proportions of the review stars and pathogenicity labels. (c) Feature importance of each classifier was assessed by SHAP values. Red indicates high, blue indicates low values of features. Model number reflects the unseen test set that was used to assess the performance. AFFIPred1 was trained on the combination of second and third variation sets and tested on the first variation set.

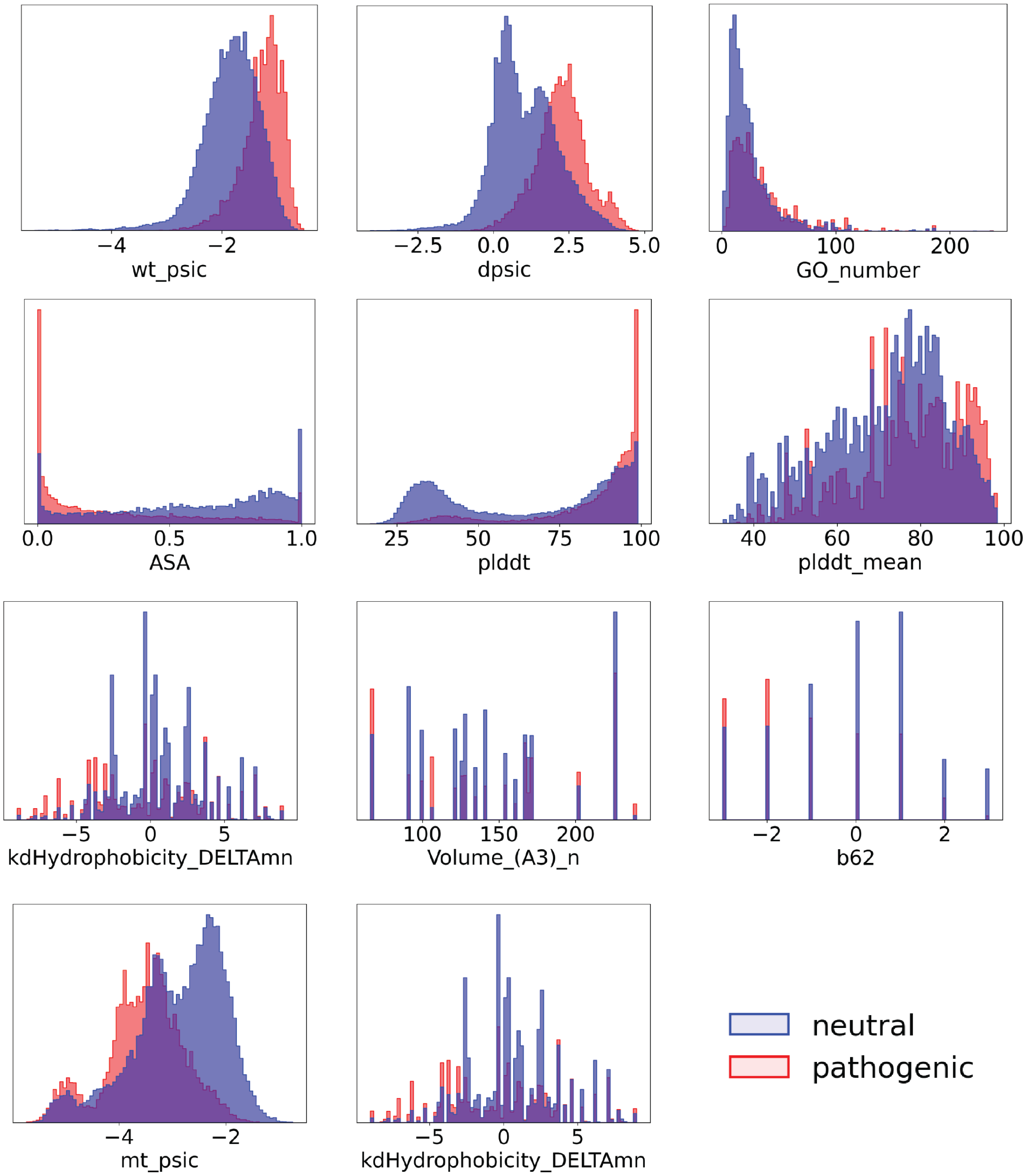


**Fig 4. Distribution of the Feature Values Across Pathogenicity Labels.**

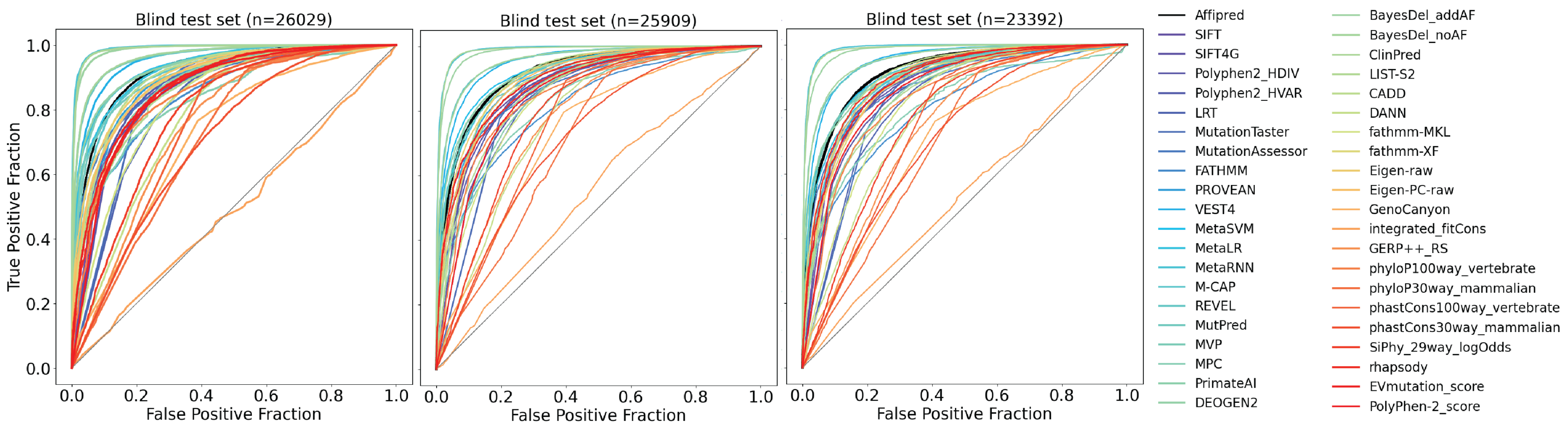


**Fig 5. AUC Results of the Benchmarking against 40 Pathogenicity Predictors.** Rank scores were used to plot ROC curves. Three panels show benchmarking of 40 tools including AFFIPred on three distinct test sets that was unseen for the AFFIPred.

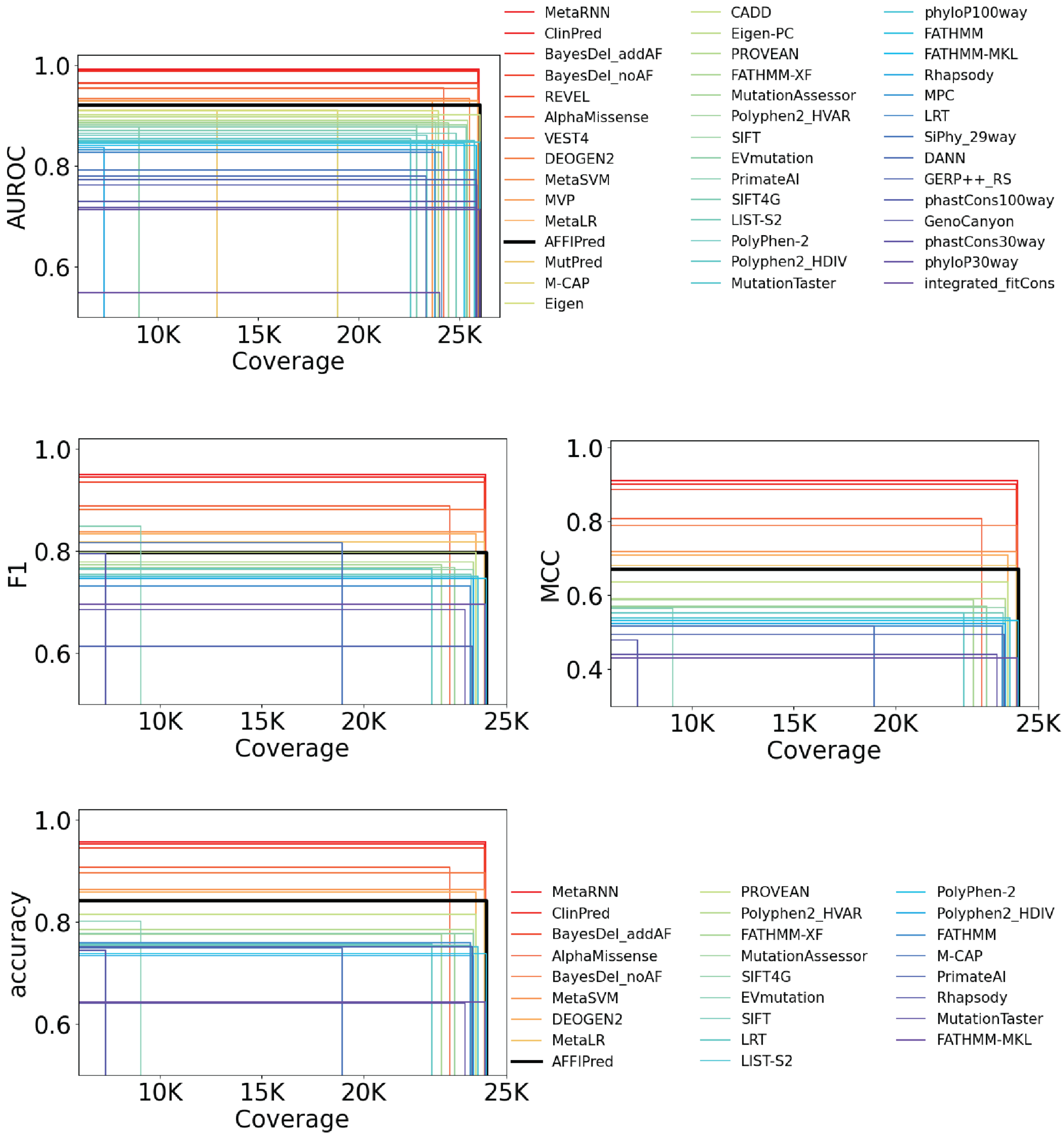


**Fig 6. Performance versus Coverage.** Plots show performance; AUC, F1, MCC or accuracy on the y-axis and the number of predicted variations on the x-axis. Three columns represent the assessments based on three distinct test sets, which was unseen for the AFFIPred and do not have any overlapping proteins.
